# Physical activity is positively related to local functional connectivity in adolescents’ brains

**DOI:** 10.1101/2020.05.29.122788

**Authors:** Ilona Ruotsalainen, Enrico Glerean, Juha Karvanen, Tetiana Gorbach, Ville Renvall, Heidi J. Syväoja, Tuija H. Tammelin, Tiina Parviainen

## Abstract

Adolescents have experienced decreased aerobic fitness levels and insufficient physical activity levels over the past decades. While both physical activity and aerobic fitness are related to physical and mental health, little is known concerning how they manifest in the brain during this stage of development, characterized by significant physical and psychosocial changes. Previous investigations have demonstrated associations of physical activity and aerobic fitness with the brain’s functional connectivity in both children and adults. However, it is difficult to generalize these results to adolescents because the development of functional connectivity has unique features during adolescence. Here, we examined how physical activity and aerobic fitness are associated with local and interhemispheric functional connectivity of the adolescent brain, as measured with resting-state functional magnetic resonance imaging (fMRI). Physical activity was measured by hip-worn accelerometers, and aerobic fitness by a maximal 20-m shuttle run test. We found that higher levels of moderate-to-vigorous physical activity, but not aerobic fitness, were linked to increased local functional connectivity as measured by regional homogeneity in 13–16-year-old participants. However, we did not find evidence for significant associations between adolescents’ physical activity or aerobic fitness and interhemispheric connectivity, as indicated by homotopic connectivity. These results suggest that physical activity, but not aerobic fitness, is related to local functional connectivity in adolescents. Moreover, physical activity shows an association with a specific brain area involved in motor functions but did not display any widespread associations with other brain regions. These results can advance our understanding of the behavior-brain associations in adolescents.

## INTRODUCTION

Although the benefits of physical activity are well recognized, the majority of adolescents are insufficiently physically active (Guthold et al., 2020; Hallal et al., 2012). This lack of physical activity has raised significant concerns, as it has been linked to the mental and physical health of adolescents (Hallal et al., 2006). Physical activity or exercise is also required to improve aerobic fitness, which is also linked to adolescents’ physical and mental health (Lang et al., 2018). A substantial global decline have been observed in the aerobic fitness levels of children and adolescents since the 1980s, and some have suggested that this decline has impacted the population health as well (Tomkinson et al., 2019). While several studies have reported that physical activity and aerobic fitness are associated with mental and physical health, little is known about their relations to brain health during the unique period of adolescence, which involves significant changes in physical characteristics, social environments, and brain properties. The present study examined the associations of both physical activity and aerobic fitness with the functional connectivity of the brain.

The functional connectivity of the brain can be measured using functional magnetic resonance imaging (fMRI). More precisely, functional connectivity measures statistical dependencies of neurophysiological events and can be used to study functional communication between or within brain regions (Friston, 2011). Studies of resting-state functional connectivity have provided important insights into the brain’s functional architecture. Moreover, functional connectivity has been shown to be associated with psychopathology in adolescents (Connolly et al., 2017; Jalbrzikowski et al., 2019; Xia et al., 2018), and is sensitive to between-subject variability e.g. symptom severity (Drysdale et al., 2017; Xia et al., 2018). Individual trait differences also outside the clinical field may manifest in functional connectivity measures, and it is therefore crucial to identify those other factors related to or influencing functional connectivity.

Previous studies of children and adults have provided evidence that physical activity and aerobic fitness may influence the functional connectivity of the brain (Boraxbekk et al., 2016; Ikuta & Loprinzi, 2019; Schaeffer et al., 2014; Talukdar et al., 2018; Tozzi et al., 2016; Voss et al., 2016). However, this relationship remains to be addressed in adolescents. It is difficult to generalize previous results concerning other age groups to adolescents because adolescence is unique period in the life-span exhibiting specific patterns of functional connectivity changes. (Váša et al., 2020).

Studies of structural connectivity have shown that physical activity and aerobic fitness are related to the white matter structure connecting the brain’s hemispheres, namely the corpus callosum. Studies in children (Chaddock-Heyman et al., 2014) and adolescents (Ruotsalainen et al., 2020) have found that aerobic fitness correlates with white matter micro-structure in the corpus callosum. Furthermore, physical activity intervention has been found to influence the corpus callosum as well (Chaddock-Heyman et al., 2018). Although functional connectivity cannot be explained solely by considering the brain’s structural connectivity, many studies have demonstrated that functional connectivity at least partly reflects underlying structural connections (Mollink et al., 2019; Shah et al., 2018; Van Den Heuvel et al., 2009). The corpus callosum has been shown to play an important role in the interhemispheric exchange of information (van der Knaap & van der Ham, 2011). Specifically, the corpus callosum has been suggested as a significant underlying structure in terms of affecting homotopic connectivity, which refers to functional connectivity among homologous brain locations across the hemispheres (De Benedictis et al., 2016; Mancuso et al., 2019; Tobyne et al., 2016). Despite these similarities, it is not yet known whether physical activity and aerobic fitness also relates to the homotopic functional connectivity of the brain.

In contrast to homotopic connectivity, which typically reflects long-range connectivity, regional homogeneity (ReHo) represents local functional connectivity in the brain. It is not yet known whether physical activity or aerobic fitness is associated with ReHo in humans. However, motor tasks have been shown to acutely modulate ReHo in sensorimotor brain areas (Lv et al., 2013). A recent study of physical exercise and the brain using animal models found that exercise induced changes in ReHo as well as in stress-related behavior in young mice with mild stress (Dong et al., 2020). These preliminary findings suggest that physical exercise influences ReHo and that these changes might be associated with factors underlying psychological wellbeing. However, this interpretation remains to be confirmed in humans. Interestingly ReHo in several brain regions has been associated with psychopathology in adolescents (Wang et al., 2018). Furthermore, some evidence suggests that ReHo can be influenced by cognitive training (Takeuchi et al., 2017) and combined cognitive-physical training (Zheng et al., 2015).

The current study aimed to examine cross-sectional associations of physical activity and aerobic fitness with the brain’s local and interhemispheric functional connectivity. First, we studied the association of both aerobic fitness (estimated using a 20-m shuttle run test) and moderate-to-vigorous physical activity (measured with accelerometers) with voxel-wise homotopic connectivity. Based on our previous findings showing an association between aerobic fitness and corpus callosum microstructure (Ruotsalainen et al., 2020), we hypothesized that aerobic fitness, but not physical activity, would be associated with 13–16-year-old adolescents’ homotopic connectivity. Further, in contrast to longer range functional connections, we studied the associations of physical activity and aerobic fitness with the brain’s local functional connectivity using ReHo. By investigating the links of physical activity and aerobic fitness with the brain’s functional connectivity, we aimed to present evidence that will allow greater insights into potential brain’s system-level connections with adolescent behavior.

## METHODS

### Participants

Participants (12.7–16.2 years old) were recruited from a larger follow-up study (Joensuu et al., 2018; Syväoja et al., 2019). A total of 61 right-handed subjects participated in the magnetic resonance imaging (MRI) scan (for participant demographics see, Table 1). One participant did not complete the resting-state functional imaging (rs-fMRI) protocol, and one participant was removed from the analysis due to excessive head motion during scanning. A total of 59 subjects (39 female) were included in the final analyses. The participants were screened for exclusion criteria comprising MRI contraindications; neurological disorders; medications influencing the central nervous system; major medical conditions; and left-handedness, which was assessed using the Edinburgh Handedness Inventory during the first research visit. Furthermore, to evaluate pubertal development, each participant was asked to self-report their stage of puberty by using the Tanner scale (Marshall & Tanner, 1969, 1970). This study was conducted according to the ethical principles stated in the Declaration of Helsinki, and each participant and his or her legal guardian provided written informed consent prior to the participation. The Central Finland Healthcare District Ethical Committee accepted this study.

**Table 1.** Participant demographics.

|  | Mean ± SD | Range |
| --- | --- | --- |
| Age (years) | 14.3 ± 0.9 | 12.7–16.2 |
| Pubertal stage (Tanner score) | 3.5 ± 0.9 | 1.5–5.0 |
| Physical activity (accelerometer, moderate-to-vigorous intensity, min/day) | 48.5 ± 19.0 | 18.0–105.9 |
| Aerobic fitness (20-m shuttle run test, min) | 5.6 ± 2.3 | 1.5–11.6 |
| BMI | 20.7 ± 2.7 | 14.6–31.1 |
| N = 59 (female = 39) |  |  |

### Physical activity and aerobic fitness

Physical activity was objectively measured using the triaxial ActiGraph GT3X+ and wGT3X+ accelerometers (Pensacola, FL, USA; for full details, see Joensuu et al., 2018). The participants were instructed to wear these devices on their right hips during waking hours for seven consecutive days (except during bathing and swimming). A valid measurement day consisted of at least 10 h of data. Subjects who had at least two valid weekdays and one valid weekend day were included in the analysis. For those subjects who did not meet these criteria, a multiple imputation method (explained in greater detail below) was employed to compensate for the missing data. Activity counts were collected in 15-s epochs. Any period of at least 30 min of consecutive zero counts was considered as a non-wear period. Data were collected at a sampling frequency of 60 Hz and standardly filtered. A customized Visual Basic macro for Excel was used for data reduction. The cut points used in the analysis were derived from a study by Evenson et al. (2008). The data regarding moderate-to-vigorous intensity physical activity was converted into a weighted-mean value of moderate-to-vigorous intensity physical activity per day ([average moderate-to-vigorous intensity physical activity min/day of weekdays × 5 + average moderate-to-vigorous intensity physical activity min/day of weekend days × 2] / 7).

A maximal 20-m shuttle run test was employed to assess the aerobic fitness of the participants. This test was performed as described by Nupponen et al. (1999) and as specified in detail for the present data collection in Joensuu et al. (2018). Each participant ran between two lines, 20 meters apart, at an accelerating pace, which was indicated with an audio signal. The duration for which each participant ran until they failed to reach the end lines within two consecutive tones indicated their level of aerobic fitness. The speed in the first and second levels were 8.0 km/h and 9.0 km/h, respectively. After the second level, the speed sequentially increased by 0.5 km/h per level. The duration of each level was one minute. The participants were verbally encouraged to keep running throughout the test. The 20-m shuttle run test is widely used to indirectly estimate aerobic fitness levels. This test was chosen for the present study because it is easily implemented with many participants, and the sub-sample that took part in the neuroimaging experiments came from a large study with nearly a thousand participants (Syväoja et al., 2019). The reliability of this test and its correlation with maximal oxygen consumption have been found to be relatively high (Castro-Pinero et al., 2010; Liu et al., 1992; Mayorga-Vega et al., 2015).

In addition to the aerobic fitness test, the participants also completed a set of tests measuring muscular fitness (push-ups and curl-ups), flexibility, and fundamental movement skills (a five-leap test and a throwing-catching combination test), which were not included in the present study.

### Multiple imputation of missing data

Multiple imputation was used to handle missing data. The proportions of missing values were 10% for the pubertal stage, 15% for the 20-m shuttle run, and 22% for the moderate-to-vigorous physical activity. Most of the missing values were due to participant absences from school during the measurements (e.g., due to sickness) and insufficient numbers of valid measurement days (i.e., less than two weekdays and one weekend day) for physical activity. Incomplete data regarding several variables were imputed using the multiple imputation under a fully conditional specification (chained equations; Van Buuren et al., 2006). The analysis was performed under the assumption of data missing at random, as the crucial predictors such as preceding measures (measured approximately six months prior to the present study) of pubertal stage, shuttle run tests (correlation with preceding 20-m shuttle run test = 0.57), and weekday measures of physical activity (correlation with the total moderate-to-vigorous physical activity [also weekend days included] = 0.95) were available. As advised (Van Buuren, 2012, chapter 2.3.3), 50 imputed datasets were constructed and analyzed. Each data set was constructed using 50 iterations of the multiple imputation by a chained equation algorithm to ensure the convergence of the iterative imputation process. The calculations were performed in R 3.4.0 (R Core Team, 2018) using the mice 2.3 package (Van Buuren & Groothuis-Oudshoorn, 2011). The model parameters and their standard errors were estimated for each imputed dataset and combined using Rubin’s rules (Van Buuren, 2012, p. 37–38) to obtain the final estimates of the parameters and their standard errors. The multiple imputation process of the present study has been described previously (Ruotsalainen et al., 2019).

### MRI acquisition

Imaging data were collected on a 3T whole-body MRI scanner (MAGNETOM Skyra, Siemens Healthcare, Erlangen, Germany) using a 32-channel head coil at the Aalto NeuroImaging unit, Aalto University, Espoo, Finland. The total scanning time was approximately 45 min for structural, diffusion-weighted, functional, field map, and perfusion imaging. All scans, except the perfusion MRI scan, were acquired using Auto Align to minimize the variations in slice positioning (van der Kouwe et al., 2005). Prior to imaging, the participants were familiarized with the measurement protocol. All participants were instructed to keep their heads still during the scanning, and pads were used to minimize head motion. In addition, earplugs were used to reduce scanner noise. During the rs-fMRI scan, participants were instructed to keep their eyes open and fixate on a cross. The rs-fMRI data was acquired using an echo-planar-imaging (EPI) sequence with the following parameters: run duration = 7 min 5 s, TR = 2610 ms, TE = 30 ms, flip angle 75°, FOV = 210 mm, 45 interleaved axial slices, GRAPPA acceleration = 2, phase partial Fourier = 7/8, and voxel size = 3.0 × 3.0 × 3.0 mm^3^. The rs-fMRI scan consisted of 160 EPI volumes.

Additionally, T1-weighted (T1w) structural magnetization-prepared rapid gradient-echo (MPRAGE) images were acquired (TI = 1100 ms, TR = 2530 ms, TE = 3.3 ms, voxel size = 1.0 × 1.0 × 1.0 mm^3^, flip angle = 7°, FOV = 256 × 256 x 176 mm^3^, and using the GRAPPA parallel imaging technique with an acceleration factor R = 2 and with 32 reference lines).

### Data preprocessing

The results included in this manuscript were derived from preprocessing performed using fMRIPrep 1.4.1 (Esteban et al., 2019), which is based on Nipype 1.2.0 (Gorgolewski et al., 2011; Gorgolewski et al., 2018). The following description of the preprocessing procedure is based on the boilerplate generated by fMRIPrep (CC0 license).

The T1w image was corrected for intensity non-uniformity with N4BiasFieldCorrection (Tustison et al., 2010), distributed with antsApplyTransforms (ANTs) 2.2.0 (Avants et al., 2008), and used as T1w reference throughout the workflow. The T1w reference was then skull-stripped with a Nipype implementation of the antsBrainExtraction.sh workflow (from ANTs) using OASIS30ANTs as the target template. Brain tissue segmentation of the cerebrospinal fluid (CSF), white-matter and gray-matter was performed on the brain-extracted T1w using FAST (Zhang et al., 2001). Volume-based spatial normalization to two standard spaces (MNI152NLin6Sym and MNI152NLin6Asym) was performed through nonlinear registration with antsRegistration (ANTs 2.2.0) using brain-extracted versions of both the T1w reference and the template. The following templates were selected for spatial normalization: the ICBM 152 non-linear 6th Generation Symmetric Average Brain Stereotaxic Registration Model (MNI152NLin6Sym), FSL’s MNI ICBM 152 non-linear 6th Generation Asymmetric Average Brain Stereotaxic Registration Model (MNI152NLin6Asym; Evans et al., 2012).

For the functional data the following preprocessing was performed. First, a reference volume and its skull-stripped version were generated using a custom methodology of fMRIPrep. The blood-oxygen-level-dependent (BOLD) reference was then co-registered to the T1w reference using FLIRT (FSL 5.0.9, Jenkinson & Smith, 2001) with the boundary-based registration (Greve & Fischl, 2009) cost-function. Co-registration was configured with nine degrees of freedom to account for distortions remaining in the BOLD reference. Head motion parameters with respect to the BOLD reference (transformation matrices, and six corresponding rotation and translation parameters) were estimated before any spatiotemporal filtering using MCFLIRT (Jenkinson et al., 2002). BOLD runs were slice-time corrected using 3dTshift from AFNI 20160207 (Cox & Hyde, 1997). The BOLD time-series (including slice-timing correction when applied) were resampled onto their original native space by applying a single, composite transform to correct for head-motion and susceptibility distortions. These resampled BOLD time series will be referred to as preprocessed BOLD in original space, or just preprocessed BOLD. The BOLD time series were resampled into several standard spaces, correspondingly generating the following spatially normalized, preprocessed BOLD runs: MNI152NLin6Sym and MNI152NLin6Asym. First, a reference volume and its skull-stripped version were generated using a custom methodology of fMRIPrep. Automatic removal of motion artifacts using independent component analysis (ICA-AROMA, Pruim et al., 2015) was performed on the pre-processed BOLD on MNI space time-series after removal of non-steady state volumes and spatial smoothing with an isotropic, Gaussian kernel of 6 mm full-width half-maximum (FWHM). Corresponding “non-aggressively” denoised runs were produced after such smoothing. Additionally, the “aggressive” noise-regressors were collected and placed in the corresponding confounds file.

Several confounding time-series were calculated based on the preprocessed BOLD: frame-wise displacement (FD), DVARS, and three region-wise global signals. FD and DVARS are calculated for each functional run, both using their implementations in Nipype (Power et al., 2014). The three global signals were extracted within the CSF, the white matter, and the whole-brain masks. The head-motion estimates calculated in the correction step were also placed within the corresponding confounds file. The confound time series derived from the head motion estimates and global signals were expanded with the inclusion of temporal derivatives and quadratic terms for each (Satterthwaite et al., 2013). Frames that exceeded a threshold of 0.5 mm FD or 2.0 standardized DVARS were annotated as motion outliers. All resamplings could be performed with a single interpolation step by composing all the pertinent transformations (i.e., head-motion transform matrices, susceptibility distortion correction when available, and co-registrations to anatomical and output spaces). Gridded (volumetric) resamplings were performed using ANTs and configured with Lanczos interpolation to minimize the smoothing effects of other kernels (Lanczos, 1964). Non-gridded (surface) resamplings were performed using mri_vol2surf (Free-Surfer).

Following fMRIPrep preprocessing, the whole-brain masked rs-fMRI data were detrended, band-pass filtered (0.008–0.08 Hz), confounds regressed, and standardized with Nilearn’s image.clean_img (version 0.2.5). The regressed confounds included six in-scanner movement estimates, the time series of the mean white matter, the mean CSF, and the mean global signal. The temporal derivatives and squares of both the original and temporal derivative time series were also included as regressors. Furthermore, a spike regressor was added for each volume for which either of the following was true: FD > 0.5 mm or DVARS > 2. This approach was previously shown to perform well in a dataset with similar motion characteristics (Parkes et al., 2018). A participant was excluded from the analysis if there were less than four minutes of their data that had FD less than 0.5 mm or DVARS < 2. One subject was removed on the basis of these criteria.

### ReHo and voxel-mirrored homotopic connectivity (VMHC)

To examine the associations of aerobic fitness and physical activity with ReHo, Kendall’s coefficient of concordance value was calculated between each voxel’s time series with its 26 neighbor voxels using the 3dReho tool from the Analysis of Functional NeuroImages (AFNI) software suite (Cox, 1996; Zang et al., 2004). Subsequently, the individual ReHo maps were standardized into z-values by subtracting the mean ReHo value throughout the whole brain and dividing it by the standard deviation. Finally, the standardized ReHo maps were spatially smoothed with a 6-mm kernel.

The voxel-mirrored homotopic connectivity (VMHC) was calculated according to the pipeline provided by the Configurable Pipeline for the Analysis of Connectomes (C-PAC; Craddock et al., 2013; Zuo et al., 2010), excluding the preprocessing steps, which were completed as described above. Before the VMHC analysis, the rs-fMRIs were spatially smoothed with a 6-mm kernel. First, the BOLD series registered onto the symmetrical anatomical template (MNI152NLin6Sym) was left-right swapped using the fslswapdim tool from FSL (Jenkinson et al., 2012). The Pearson correlation was then calculated for each voxel and its mirrored counterpart. Further, the correlation values were Fisher z-transformed, and the participant-specific Z statistic maps were used in the statistical analysis.

To test the associations of physical activity and aerobic fitness with ReHo and VMHC, we used a FSL’s randomise tool with a nonparametric permutation test with 10,000 permutations and variance smoothing (Winkler et al., 2014). Age, pubertal stage, and sex were used as covariates. The T-value difference in the voxel clusters was considered noteworthy when the values passed – after threshold-free cluster enhancement (TFCE) and family-wise error correction – a threshold of cluster corrected p < 0.05. The cluster that passed these criteria was labelled according to the Harvard-Oxford Cortical Structural Atlas (Desikan et al., 2006; Frazier et al., 2005; Goldstein et al., 2007; Makris et al., 2006). The code for the ReHo and VMHC analyses is available at https://osf.io/ydwhz/.

## RESULTS

### Physical activity, aerobic fitness, and ReHo

In the present study, we sought to understand how physical activity and aerobic fitness are related to ReHo in 13–16-year-old adolescents. We analyzed data from 59 participants using FSL’s randomise with nonparametric permutation tests taking into account age, sex, and pubertal stage. After TFCE and family-wise error correction, we found that higher moderate-to-vigorous physical activity was correlated with increased ReHo in one cluster located mainly in the right supramarginal gyrus (see Fig. 1 and Table 2 for cluster details). Concerning aerobic fitness, we did not find any noteworthy correlations with ReHo (see Fig. 2 for non-threshold associations of ReHo with physical activity and aerobic fitness). As head motion may have affected the results, we also conducted an analysis with average FD as an additional regressor; this had negligible effects on the results (Supplementary Fig. 1). All statistical maps can be found in our Neurovault collection at neurovault.org/collections/QPRRARMZ/ (Gorgolewski et al., 2015).

**Table 1.** A significant positive correlation between physical activity and ReHo. The MNI coordinates specify the location of the center of mass and the p-value is corrected for familywise error and TFCE. All tracts overlapping that the cluster are labeled according to the Harvard-Oxford Cortical Structural Atlas. The proportion of voxels overlapping that particular brain area is shown in parentheses.

| Cluster size<br>(voxels) | Structures to which the cluster belongs to (% of voxels) | p-value | MNI coordinates |  |  |
| --- | --- | --- | --- | --- | --- |
|  |  |  | X | Y | Z |
| 33 | Supramarginal gyrus (51.8), planum temporale (10.0), superior temporal gyrus (7.8), parietal operculum cortex (6.6), angular gyrus (0.2) | 0.035 | 65.2 | -32 | 30.8 |

**Figure 1.**
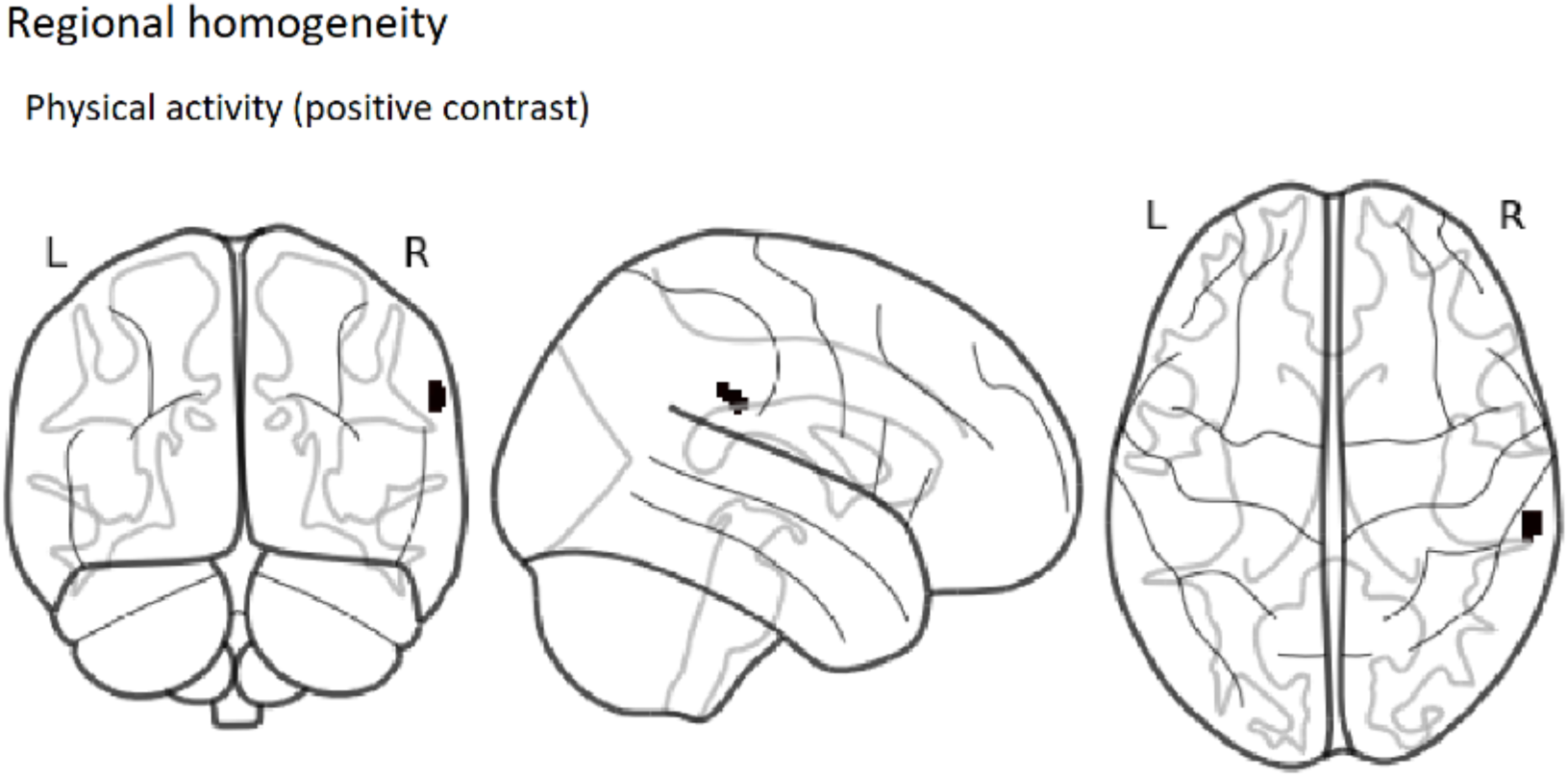
Group-level thresholded map of positive contrast (correlation) between moderate-to-vigorous physical activity and ReHo (p < 0.05, corrected for TFCE and family-wise error). The colored area represents the region significantly correlated with physical activity.

**Figure 2.**
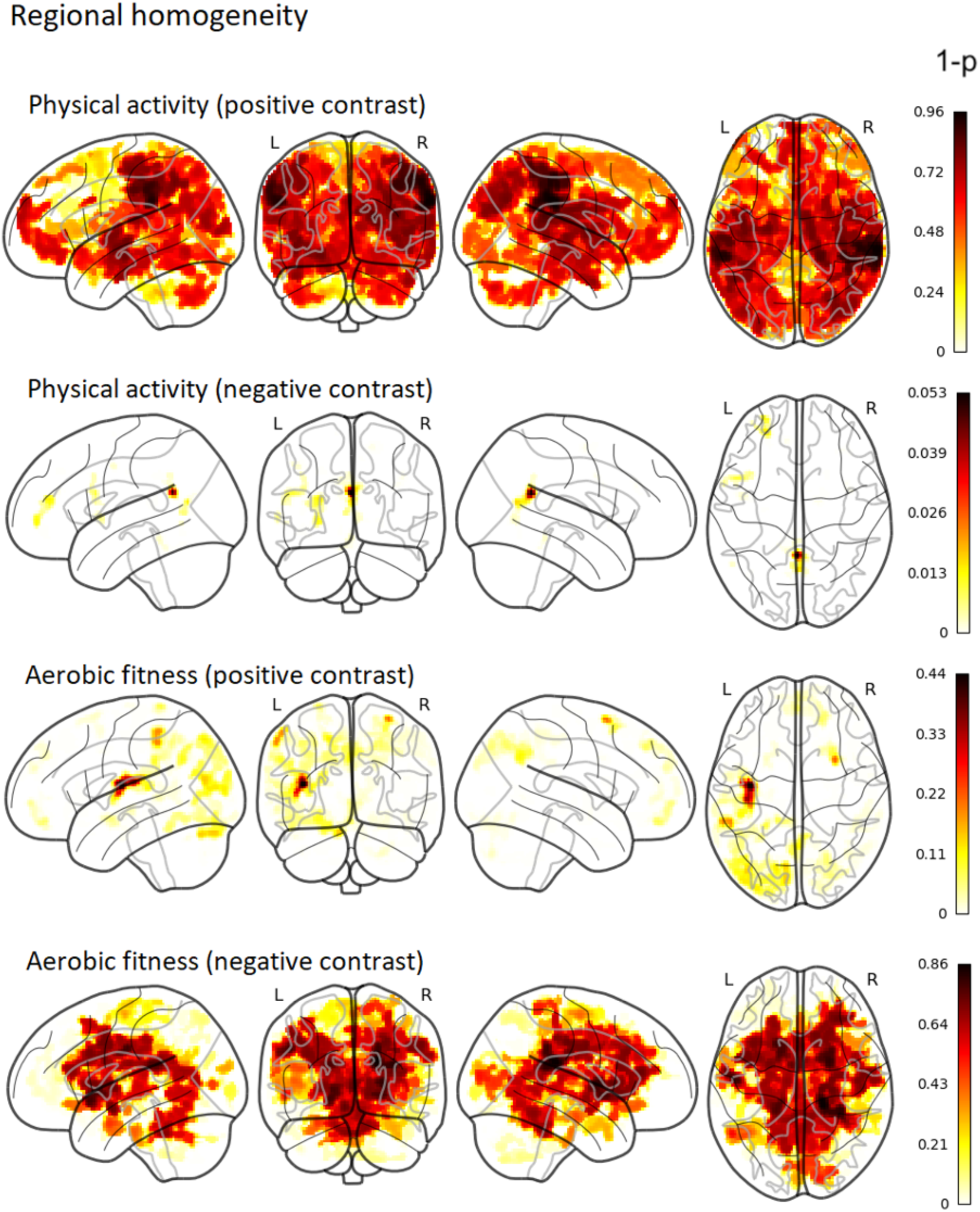
Group-level 1-p-value maps (non-thresholded) of positive and negative contrasts (correlations) of physical activity and aerobic fitness with ReHo (corrected for TFCE and family-wise error). Intensity of color represents 1-p-value (both significant and nonsignificant associations are shown). Thus, the darker color (red) represents a smaller p-value, whereas the lighter color (yellow) represents a larger p-value.

### Physical activity, aerobic fitness, and homotopic connectivity

VMHC was used to study the correlations of physical activity and aerobic fitness with homologous interhemispheric connectivity. After TFCE and family-wise error correction, neither physical activity nor aerobic fitness showed any noteworthy correlation with homotopic connectivity (Fig. 3). All statistical maps can be found in our Neurovault collection at neurovault.org/collections/QPRRARMZ/ (Gorgolewski et al., 2015).

**Figure 3.**
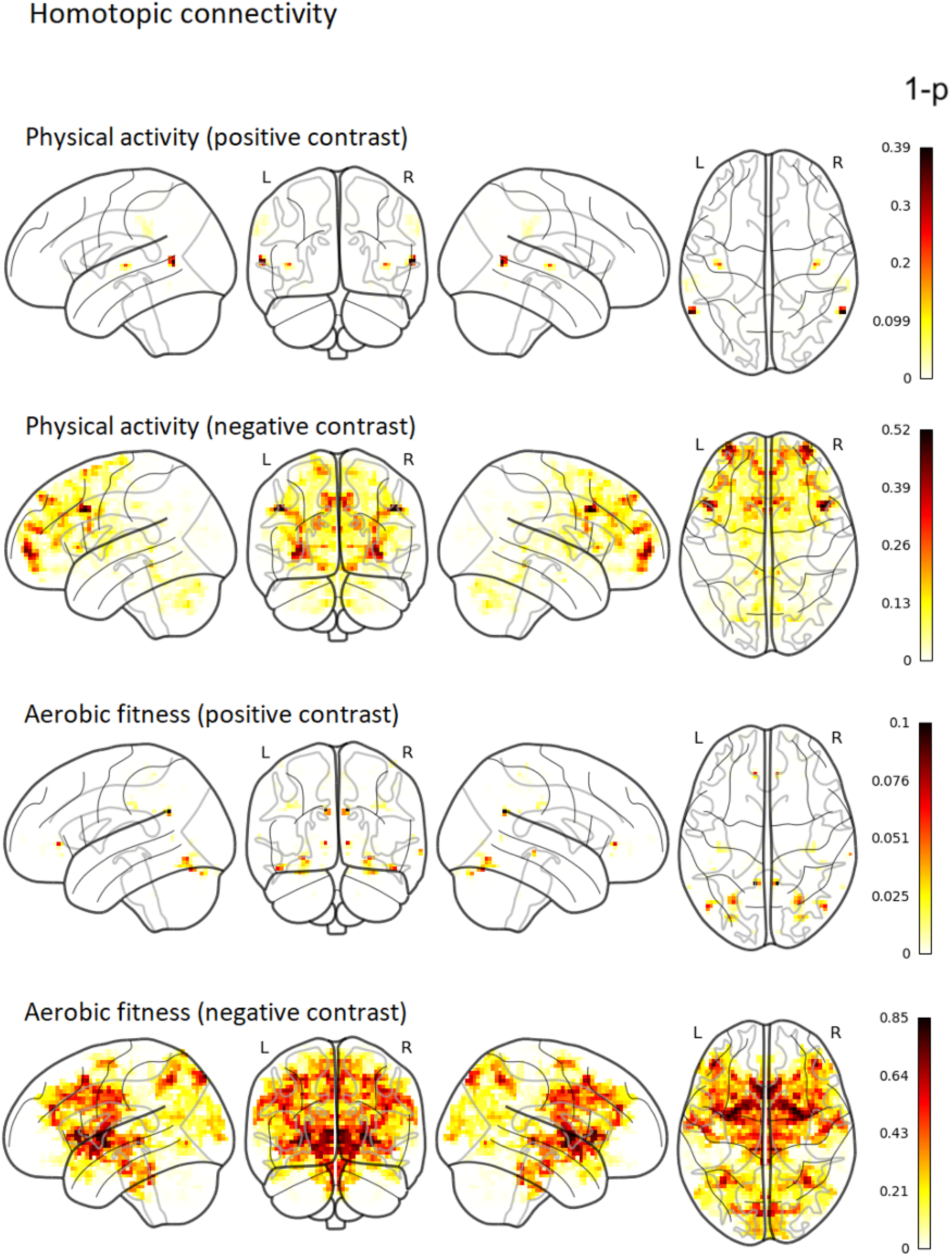
Group-level non-thresholded 1-p-value maps for the nonsignificant associations of physical activity and aerobic fitness with VMHC (corrected for TFCE and family-wise error). Intensity of color represents 1-p-value (both significant and nonsignificant associations are shown). Thus, the darker color (red) represents a smaller p-value, whereas the lighter color (yellow) represents a larger p-value.

## DISCUSSION

We explored whether physical activity and aerobic fitness levels are associated with the brain’s functional connectivity in adolescents, whose brains, bodies, and sociopsychological environments undergo significant changes within a short time period. More specifically, we examined whether moderate-to-vigorous physical activity and aerobic fitness are associated with local and interhemispheric functional connectivity, as indicated by ReHo and VMHC, respectively. The resulting data revealed a positive association between physical activity and ReHo in a cluster located mainly in the right supramarginal gyrus. However, we did not find a significant association between aerobic fitness and ReHo. Contrary to our expectations, we did not find evidence for an association between homotopic connectivity and aerobic fitness or with physical activity. Overall, these findings suggest a link between adolescents’ physical behavior and the local functional connectivity of their brains.

The positive association between physical activity and ReHo was located mainly in the right supramarginal gyrus. This area is classically defined as a part of the somatosensory cortex, but has since been associated with many higher cognitive functions, including language (Oberhuber et al., 2016; Stoeckel et al., 2009) and empathy (Silani et al., 2013). More importantly in the context of the present findings, the supramarginal gyrus is also involved in proprioception (Ben-Shabat et al., 2015; Kheradmand et al., 2015) as well as in motor attention and planning (Barbaro et al., 2019; Burke et al., 2013; McDowell et al., 2018; Rushworth et al., 1997). Proprioception refers to a sense of movement and body position (Tuthill & Azim, 2018). To engage in physical activity, it is crucial to be able to sense one’s own body and movement. The ability to plan movements and maintain a motor plan is also critical to physical activity. Our results could therefore indicate that physical activity increases local functional connectivity in specific motor-related areas of the adolescent brain.

However, intervention studies are needed to confirm this causal relationship, as the opposite interpretation of this association is also possible.

Interestingly, an earlier study found that executing a hand-closing-opening task motor task acutely decreased ReHo in the right supra-marginal gyrus relative to a resting state (Deng et al., 2016). Changes in ReHo have also been reported in relation to a finger tapping task (Lv et al., 2013). Specifically, Lv et al. (2013) showed that finger-tapping speed modulated the ReHo response in the sensorimotor cortex. While the execution of slow finger-tapping movement was associated with greater ReHo relative to a resting state, fast movement was associated with lower ReHo relative to a resting state, when measured at commonly used low frequency band (0–0.08 Hz). Overall, it appears that movement execution relates to local functional communication measured with ReHo. The aforementioned studies demonstrated an association between acute changes in local functional connectivity and sensorimotor behavior. However, they did not reveal the long-term effects of continued physical performance on local functional connectivity.

There is a lack of studies investigating the association of physical activity and aerobic fitness with ReHo in humans. However, additional evidence relating to this issue has been gained through animal experiments. The present results are partly in line with those of previous studies of young mice. Dong et al. (2020) showed that physical exercise (wheel running) caused widespread increases in ReHo and decreases in stress-related behavior in young mice with mild stress. In the present study, we did not find widespread associations between physical activity and ReHo, but we did find such an association in a small cluster in a region important for motor actions and/or movement sensations. ReHo changes in specific brain regions have been suggested as potential neuroimaging markers of mental health problems in adolescents (Wang et al., 2018), and physical activity relates to mental health.

However, the present findings suggest it is unlikely that ReHo could be the mechanism underlying the positive effect of physical activity on mental health in adolescents, as the association was limited to a small area. However, this conclusion remains to be confirmed by future studies.

Despite the fact that ReHo was associated with physical activity, we did not observe any noteworthy connections to aerobic fitness. By measuring physical activity, we measure movement produced by muscles requiring energy expenditure (Caspersen et al., 1985). Aerobic or cardiorespiratory fitness, on the other hand, refers to the body’s ability to deliver oxygen to the muscles during sustained physical activity or exercise (Caspersen et al., 1985). Thus, aerobic fitness refers to a capacity that a person has obtained over previous months or years, partly affected by heredity. Physical activity, herein measured with accelerometers over a seven-day period, refers to the current physical behavior of a person. The connection of ReHo with physical activity but not with aerobic fitness could mean that, in the context of physical performance, behavior is a more important determinant of ReHo changes in the brain compared to capacity. This interpretation is in line with earlier findings, which demonstrated that motor behavior (i.e., physical exercise and motor skill learning) increases local functional connectivity and autonomy in the sensorimotor areas over the course of several weeks (Bassett et al., 2015; Dong et al., 2020). In light of these earlier findings showing that motor behavior can influence local functional connectivity, our result may reflect the adaptation of the supramarginal gyrus to physical activity.

Concerning interhemispheric homotopic connectivity, contrary to our expectations, we did not find any noteworthy associations with either physical activity or aerobic fitness. In our previous study of the same sample of adolescents, we found that aerobic fitness correlates with white matter properties of the corpus callosum (Ruotsalainen et al., 2020). The corpus callosum plays a crucial role in interhemispheric functional connectivity, as it is the main white matter structure that connects the brain’s hemispheres. Indeed, the properties of the corpus callosum are significantly related to homotopic connectivity (De Benedictis et al., 2016; Mancuso et al., 2019; Tobyne et al., 2016). Despite the demonstrated link between aerobic fitness and the corpus callosum, we did not find a clear evidence of a connection between aerobic fitness and homotopic connectivity. This finding might be explained by the fact that even though the corpus callosum influences homotopic connectivity, its role appears to be small. Mollink et al. (2019) found that white matter microstructure explains 1–13 % of a variance in homotopic connectivity. Interestingly, the relationship between structural and functional connectivity increases with age (Betzel et al., 2014). Therefore, it is possible that the connection of these two factors is not as prominent in adolescents as in adults.

Previous literature concerning the associations of between physical activity and aerobic fitness with interhemispheric functional connectivity in healthy participants is scarce. In a study of healthy older adults, physical activity did not relate to interhemispheric connectivity (Veldsman et al., 2017). In a study of middle-aged adults, on the other hand, Ikuta & Loprinzi (2019) found that aerobic fitness was associated with homotopic parahippocampal functional connectivity, but not with hippocampal connectivity. While our results are in line with those of Veldsman et al. (2017), some methodological and sample-related differences between the two studies should be noted. First, the present study focused on the functional connectivity of homologous brain areas (i.e., homotopic connectivity), whereas Veldsman et al. (2017) compared interhemispheric connectivity between the seed region and the brain network. Furthermore, the age range of the participants differed significantly between the two studies. Nevertheless, the findings of both studies indicated a lack of influence of physical activity on interhemispheric functional connectivity. Concerning aerobic fitness, we could not replicate the findings of Ikuta & Loprinzi (2019) with regard to the association with homotopic parahippocampal connectivity in the adolescent population. This might be explained by the larger sample size of the earlier study or the possibility that this connection with fitness is more prominent in adulthood than in adolescence.

The present study has some limitations that should be noted. First, it is recommended that all imputed data should be used in the analysis, and it is also recommended the parameters and their standard errors should be combined using Rubin’s rules (Van Buuren, 2012, p. 37– 38). However, due to the computational restrictions of neuroimaging analysis, we used averages of the imputed values in our analyses. Second, we estimated the participants’ levels of aerobic fitness using a maximal 20-m shuttle run test, which is an indirect measure of cardiorespiratory fitness. Previous studies suggest that the reliability of the maximal 20-m shuttle run test and its correlation with maximal oxygen consumption are relatively high when compared to those of direct measurements (Castro-Pinero et al., 2010; Liu et al., 1992; Mayorga-Vega et al., 2015; but see Armostrong & Welsman, 2019 for concerns regarding validity). Third, the use of different cut-points to determine the participants’ levels of moderate and vigorous physical activity might results in different estimates of the amount of moderate-to-vigorous physical activity. This could be a limiting factor for the comparison of results between similar studies. Finally, we did not include body mass index (BMI) as an additional covariate in the present analysis. BMI is related to performance in the 20-m shuttle run task. However, it does not have a large influence on the correlation between task performance and maximal oxygen consumption (Mahar et al., 2018). The present sample included only one obese participant, and excluding this participant from the analysis did not affect the results.

## CONCLUSIONS

The functional connectivity of the brain has been suggested as an important contributor to individual variation, for example, in context of adolescent mental health. The identification of factors related to functional connectivity during this period, in which many unique changes occur in the brain, can advance our understanding of the behavior-brain relationship in adolescents. We found that higher levels of accelerometer-measured moderate-to-vigorous physical activity were linked to increased local functional connectivity in 13–16-year-old adolescents. However, we did not find a clear association between aerobic fitness and local functional connectivity. Contrary to our hypothesis, neither aerobic fitness nor physical activity showed any significant correlation with interhemispheric homotopic connectivity. Our results did show that physical behavior is related to local functional connectivity. However, we did not find any strong evidence for the relationship between adolescent’s physical capacity (aerobic fitness) and local functional connectivity. Moreover, according to the present findings, physical activity appears to be related to a specific brain area involved in sensorimotor functions but does not appear to be related to widespread brain regions.

## Acknowledgments

Funding for this research was provided by the Academy of Finland [grant numbers 273971, 274086 and 311877], the Alfred Kordelin Foundation, and the Emil Aaltonen Foundation. We thank Marita Kattelus, Riikka Pasanen, and Jenni Silvo for their valuable help in the data collection. We also would like to thank Dr. Toni Auranen and Prof. Veikko Jousmäki for providing the research infrastructure for this work.

---

**Supplementary Figure 1.**
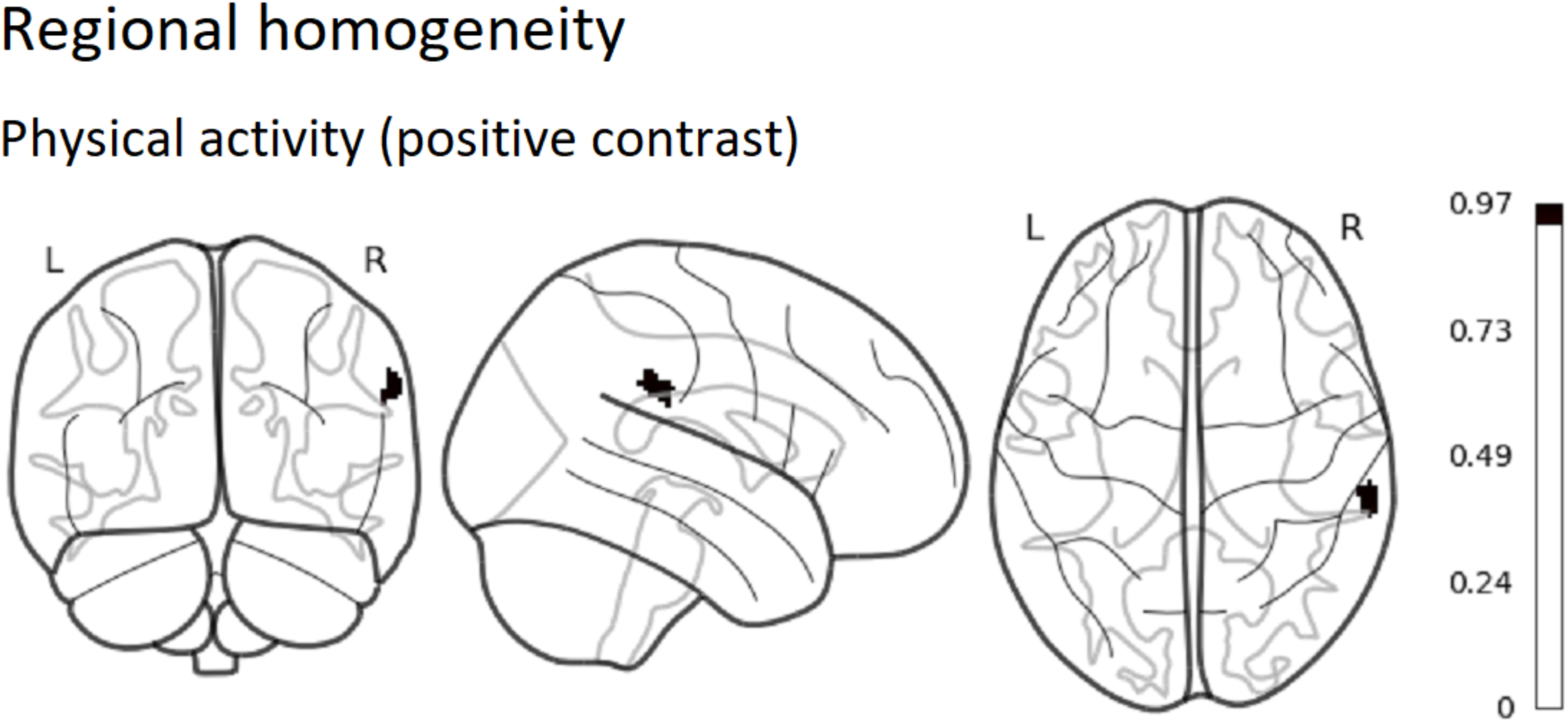
Group-level 1-p-value map regarding positive correlation between moderate-to-vigorous physical activity and ReHo (p<0.05, corrected for TFCE and family-wise error rate), when taking into account mean frame-wise displacement.

## Notes

https://neurovault.org/collections/QPRRARMZ/

### Competing Interest Statement

The authors have declared no competing interest.

